# Endemic and invasion dynamics of wild tomato species on the Galápagos Islands, across two centuries of collection records

**DOI:** 10.1101/2025.01.09.632230

**Authors:** Alex D. Kutza, Zoe L. Hert, Leonie C. Moyle

## Abstract

- We aggregated digitized herbarium and other collection records—spanning >225 years since 1795—to assess the biological, geographical, and historical factors shaping distributions of three wild tomato species on the Galápagos Islands, and to infer future threats to the two endemic species (*Solanum cheesmaniae* and *S. galapagense*) and risks posed by their invasive congener (*S. pimpinellifolium*).
- Combining >400 unique geolocated Galápagos records with bioclimate data and species distribution modelling, we quantified the geo-spatial distribution of each species, bracketed the historical timing and location of introductions of the invasive species, characterized species bioclimate envelopes, and projected suitable habitat overlap.
- We infer that dispersal limitation and alternative selective histories shape current species distributions, and that anthropogenic change has and will continue to have different impacts on the two endemic species—closely associated with their different geographic and environmental distributions. We also identify plausible avenues for, and limits to, future invasive expansion.
- These data vastly extend the temporal and spatial reach of our direct historical inferences, provide a critical complement to genomic analyses of contemporary Galápagos populations, and demonstrate that scientific collections are especially valuable for interpreting factors shaping species distributions on high-endemism islands with recent rapid environmental change.

## Introduction

Anthropogenic change is rapidly altering the distribution and diversity of endemic organisms (Manes et al., 2021), effects that can be exacerbated by interactions with invasive species (Powell et al., 2011). However, the factors that drive these changes are often poorly understood, partly because many studies are limited to recent timescales. Historical collections can often provide this missing temporal dimension (Meineke et al., 2018). By extending the representation of species sampling up to hundreds of years into the past, these data integrate over timescales more reflective of both ecological and evolutionary processes. Simultaneously, historical collections can substantially expand the known occurrence locations of individual species. This enriched spatial data can strengthen inferences about the ecological factors defining species distributions, and the nature and magnitude of differences between species. When examined over time, these records can also bracket key timepoints and critical changes in environmental factors that might alter species abundance and distribution. Together, these data are essential for understanding the temporal dynamics of individual species, and their interactions within communities, especially in sensitive locations with particular vulnerability to anthropogenic change.

Island systems are among these especially sensitive communities (Caujape-Castells et al., 2011). Ocean island and archipelagos are uniquely vulnerable to rapid environmental change from both local (e.g. land-use) and global (e.g. climate change) sources (Fordham and Brook, 2010; Harter et al., 2015; Braje et al., 2017). Moreover, greater than 31% of the world’s endemic plant species are found exclusively on islands (Schrader et al., 2024). These features make herbarium collections from islands a valuable scientific resource. First, because many island communities have experienced recent rapid change, especially over the last century (e.g., Watson et al., 2010; Escobar-Camacho et al., 2021), herbarium collections offer a record of past conditions that might differ radically from contemporary circumstances. Second, because field surveys can be especially challenging in remote or inaccessible locations, like island archipelagos, herbarium specimens provide an enriched spatial complement to contemporary assessments. Herbaria are therefore a unique resource for assessing the history of island plant species, the patterns and causes of change over recent time in island communities, and the projection of these changes into the future.

Fortunately, their characteristic endemism means that islands have also been a focus of scientific collections. The Galápagos Archipelago is no exception in this regard. Although small in landmass area (<8000km2), remote (960km) from the nearest continent of South America (Escobar-Camacho et al., 2021), and only permanently colonized by humans in the 1832 (Perry, 1984), these islands have attracted scientific interest for more than two centuries. Archibald Menzies, a Scottish naturalist, collected the first known botanical specimens from the Galápagos in 1795 (Nelson and Porter, 2011). Of his three specimens, now housed at the Natural History Museum (NHM) London, one is *Solanum galapagense*—an endemic wild tomato species, collected on Isabela (Figure 1) (Nelson and Porter, 2011). In January 1825, John Scouler and David Douglas, former classmates at the University of Glasgow, Scotland, spent two days collecting zoological and botanical specimens on Santiago Island (Scouler, 1905; Robinson, 1902), including another *S. galapagense*. After the H.M.S. Beagle expedition landed on the archipelago in September 1835 (Darwin, 1845), Charles Darwin amassed 173 plant collections from four islands over the subsequent five weeks (Egerton 2010). These collections included the type specimen of *Solanum cheesmaniae*—the sister species to *S. galapagense*— which is currently held in the Cambridge University Herbarium. The Galápagos islands were also the target of dedicated scientific collecting trips in the late 19^th^/early 20^th^ century, most notably the Hopkins-Stanford and California Academy scientific expeditions in 1898/1899 and 1905/1906, respectively (Robinson, 1902; Stewart, 1911).

**Figure 1.**
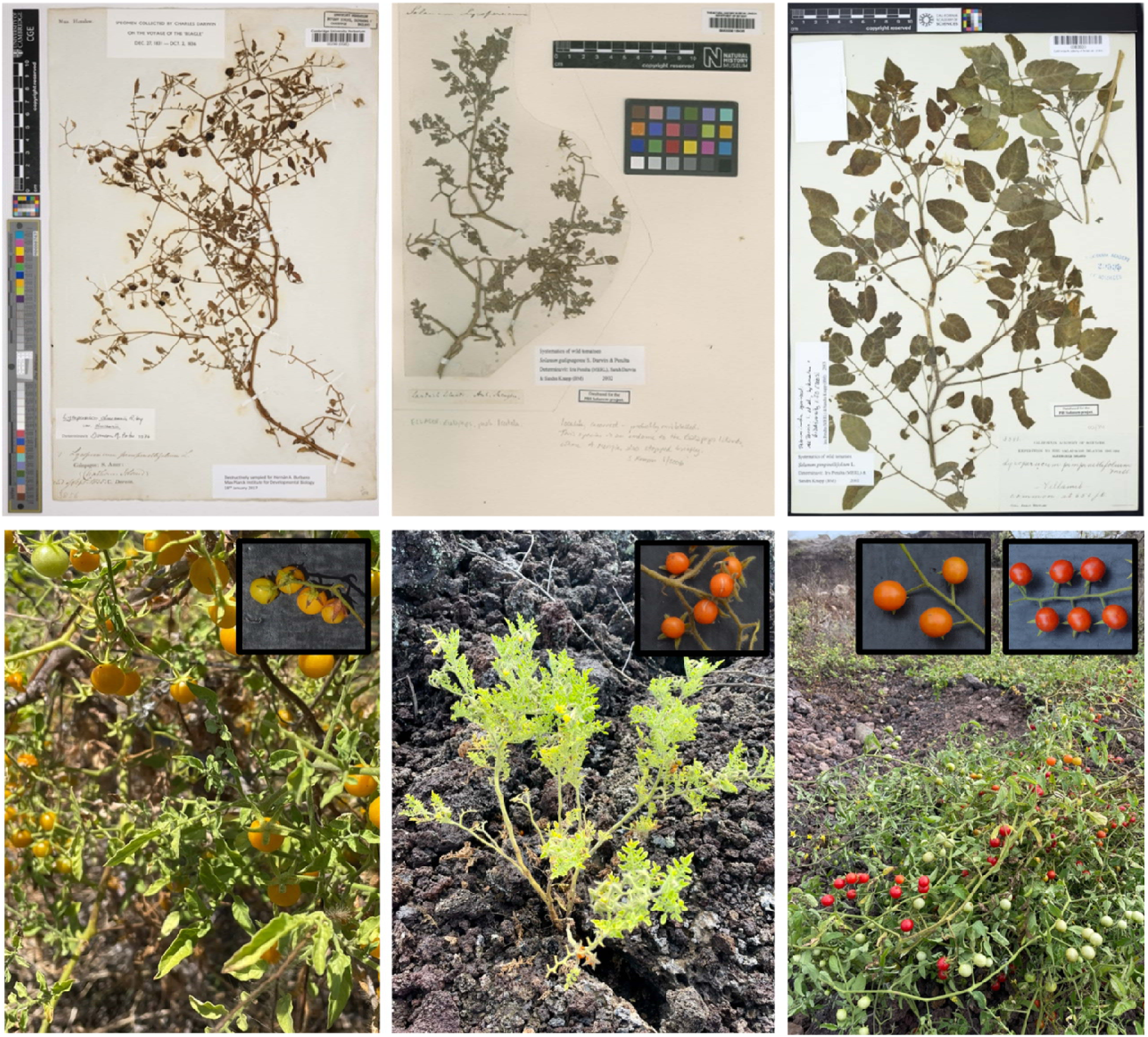
Wild tomato species on the Galápagos Islands. Upper panels: Notable herbarium specimens. (Left) Charles Darwin’s 1835 collection of *S. cheesmaniae* from San Cristóbal (‘Chatham’); CC license image courtesy of Cambridge University Herbarium (barcode: CGE00000298). (Center) Archibald Menzies’ 1795 collection of *S. galapagense* from Isabela Island; CC license image courtesy of The Trustees of the Natural History Museum, London. (Right) Alban Stewart’s 1905 collection, putatively *S. pimpinellifolium*, from Isabela; CC license image courtesy of California Academy of Sciences. **Lower panels: Contemporary observations**. (Left) *Solanum cheesmaniae*, (Center) *S. galapagense*, and (Right) *S. pimpinellifolium*, *in situ* on the Galápagos Islands, with respective fruits (insets), not to scale. (Images: L. C. Moyle).

Since these historical collections, the Galápagos Archipelago has experienced substantial changes, from both altered global climate (Wolf, 2010; Lui et al., 2013; Paltan et al., 2021) and focused waves of anthropogenic environmental change (Watson et al., 2010; Giefer, 2023).

Important terrestrial elements of the latter include: the introduction and rapid expansion of alien herbivores (e.g., goats, pigs, donkeys) in the 19^th^ and early 20^th^ century, and their subsequent concerted eradication from specific islands (Cruz et al., 2005; Carrion et al., 2007; Carrion et al., 2011; Phillips et al., 2012); the 20^th^ century growth of resident human populations (to ∼30,000 at present), in conjunction the expansion of agriculture (Giefer, 2023); and the accelerating expansion of tourism. Between 2000 and 2019, tourist numbers grew from 65,000 to > 270,000 annually (Escobar-Camacho et al., 2021). This rapid increase in the human population—both resident and tourist—over recent time has specifically been accompanied by increases in the arrival and dispersal of alien species (Toral-Granda et al., 2017). The downstream effects of these diverse human impacts—especially habitat alteration from land development, the introduction of nonnative herbivores, and competition from invasive plants—are all known to alter the distribution and genetic diversity of endemic plants on the Galápagos (Ellis-Soto et al., 2017; Torres et al., 2023). Among the species potentially affected by these changes are the two Galápagos endemic wild tomato species *S. cheesmaniae* and *S. galapagense* (Figure 1). In previous work with contemporary populations of these two endemics, we used fieldwork and population genomics to infer evidence for contraction of endemic populations (especially in proximity to human settlements); simultaneously, following others (Darwin et al., 2003; Nuez et al., 2004), we have documented the geographical expansion of a third wild tomato species— *Solanum pimpinellifolium*—recently introduced from mainland Ecuador (Gibson et al., 2020, and see ‘Study system’). Our genomic data confirms field assessments that there is recent and ongoing hybridization between endemic and invasive species (Gibson et al., 2020; Gibson et al., 2021). However, because these contemporary samples are limited in time and space, our assessment of the timing and generality of these demographic and evolutionary changes is both indirect and incomplete.

Here our goal was to use digitized herbarium and collection records of these three species as a unique complement to our contemporary data. With these geospatial and temporal records, in conjunction with climate and environmental data, we: 1) quantified the geo-spatial distribution of each species on the archipelago, and bracketed the historical timing, location, and number of introductions of the invasive species; 2) characterized species differences in habitat envelopes; 3) evaluated risks to the two endemic species, from projected drivers of environmental change on the Galápagos in general, and specifically from contact with their introduced congener; and, inferred the potential locations of further expansion of the invasive species, including potential environmental limits to this expansion. By extending the temporal and spatial reach of our historical inferences, this dataset allows us to identify critical biological differences between species, and to assess past demographic changes and project future ones, for these three charismatic Galápagos plant species.

## Materials and Methods

### Study system

Our analyses focus on three species within the wild tomato group *Solanum* section *Lycopersicum*, a closely-related clade of 12 species within the >1200 species of the hyperdiverse genus Solanum (Echeverría-Londoño et al., 2020). Of these three lineages, *S. cheesmaniae* and *S. galapagense* are sister species, endemic to the Galápagos Islands (Darwin et al., 2003) and diverged from each other for as little as several thousand generations (Gibson et al. 2021) and up to 250 kya (Pease et al. 2016). Taxonomic naming of these endemic lineages has changed numerous times over the last 200 years (e.g., Stewart, 1911; Muller, 1940; Luckwill, 1943; Rick, 1956), prior to their formal classification as two separate species in 2003 (Darwin et al., 2003). This dynamism means that herbarium labels, and sometimes database records, may reflect older taxonomic conventions, occasionally resulting in challenges for parsing species identity, as we further expand below.

Although both Galápagos endemic tomato species are currently classified as “least concern” by the International Union for Conservation of Nature (IUCN), recent findings indicate they could be threatened by ongoing environmental change. In 2018-2019 field work, we found high frequencies of endemic population extirpation (more than 80% of *S. cheesmaniae* and *S. galapagense* populations previously described in 2000-2003 could not be relocated) and inferred that *S. cheesmaniae* is especially rare on islands with anthropogenic disturbance (Gibson et al., 2020). *S. cheesmaniae* also faces new threats from a third wild species, *S. pimpinellifolium*. This wild species’ natural range spans low to mid-elevation mainland South America, from Columbia to Peru (Zuriaga et al., 2009; Nakazato et al., 2010; Gibson and Moyle 2020), but it is now found on the Galápagos Islands of Santa Cruz, San Cristóbal, and Isabela (Gibson et al. 2020). The scientific literature first reported relatively large populations of this species in 1995 to 2000 (Darwin et al., 2003; Nuez et al., 2004; Darwin, 2009), primarily on Santa Cruz but secondarily (in 2000) on San Cristóbal and Isabela (Darwin et al., 2003).

However, as noted in Darwin (2009), evidence of *S. pimpinellifolium* appears in field guides as early as 1983, suggesting a notable presence on Santa Cruz by at least this date. Using population genomics, we inferred that *S. pimpinellifolium* was introduced within the last 200 generations (Gibson et al., 2021), consistent with human-mediated dispersal. Most contemporary invasive populations descend from a single introduction from mid-elevation Ecuador, although genetic ancestry indicates at least two other independent, and more recent, introductions have also occurred—one each from Ecuador and Peru (Gibson et al., 2021). In addition, we infer at least one back-migration of *S. pimpinellifolium* from the Galápagos to mainland Ecuador—represented by a mainland accession collected in Los Ríos, Ecuador (LA0411; tgrc.ucdavis.edu) (Gibson et al. 2021). As this collection dates from 1956, this implies a presence on the Galápagos since at least 1956. On Santa Cruz, contemporary field and genomic data indicate that *S. pimpinellifolium* cooccurs and hybridizes with *S. cheesmaniae*; F1 and F2 individuals have been documented at current contact sites, and genome-wide ancestry inference indicates a hybridization history of up to 12 generations (Gibson et al., 2021). In the present study, we aimed to use digitized herbarium and collections data to better understand the biological significance of these contemporary observations, now and into the future.

### Data collection

#### A. Herbarium records

Our initial list of herbarium records was obtained by querying the Solanum Planetary Biodiversity Inventory (PBI) database (https://solanaceaesource.myspecies.info/) (with the kind assistance of S. Knapp; NHM), and retaining records of our three target species (*Solanum cheesmaniae, Solanum galapagense*, and *Solanum pimpinellifolium*) whose geolocation was the Galápagos Islands. We also obtained herbarium and/or geolocated sample records by directly querying the California Academy of Science herbarium collections (CAS), the U.C. Davis Center for Plant Diversity Herbarium (DAV), and the U.C. Davis C. M. Rick Tomato Genetics Resource Center (TGRC). These three sources represent a substantial fraction of the 20^th^ century herbarium and other collections of Galápagos Solanum (see *Results*). Note that the TGRC data are not herbarium records, but geolocation and associated data from germplasm collections. In total, queries returned >800 herbarium or other observation/collection records.

From this initial list, we took several steps to cross-validate and/or remove redundant records, and to ensure recorded species names reflect contemporary taxonomic classifications (as in Darwin et al., 2003). First, for each of the PBI records, we visited the individual online herbarium databases to confirm that these aligned with available digital data and/or specimen records. We were unable to directly cross validate 150 of these records, either because the ‘home’ herbarium did not have publicly available data (N=127), or because the online databases were inaccessible due to maintenance, or publicly available data did not include the specific record (N=15 cases, from 7 different herbaria). These records were retained in our complete list under the assumption that the PBI has curated a more expansive dataset, as most of these records were from the Charles Darwin Research Station (CDS). Second, across our whole dataset we removed records of specimens that were duplicates of the same time, location, and species, to ensure only one unique record was included for downstream analyses. (This was most apparent with duplicate specimens present in DAV and the TGRC, in which case we eliminated the TGRC duplicate record.) In cases where specimens with identical date, location, and species were recorded in two herbaria, and it was unclear whether the physical specimen(s) existed in either or both locations (N=9), we retained the record associated with the herbarium that has the greatest number of other Solanum Galápagos specimens. In every case, we have annotated the verification status of every record in our dataset, including whether it was listed in multiple herbaria, whether it was directly verified, and whether we have adjusted the species name to reflect contemporary nomenclature.

In addition, to identify or confirm additional older (pre-20^th^ century) collection records, we queried several individual herbarium databases, including the Cambridge University Herbarium (‘CGE’), the Natural History Museum (NHM) London (‘BM’), and the Royal Botanic Garden Edinburgh (‘E’) (see Table S1 for all pre-20^th^ century records). CGE’s records of Darwin’s collections included four from the Galápagos: three *Solanum cheesmaniae* specimens (including an isolectotype) and a *Solanum galapagense* specimen. NHM London houses the earliest known *S. galapagense* specimen, from 1795 collected by Archibald Menzies. (Note: the collection date in the NMH database is 1791, however Menzies expedition to the Galápagos was 1795 (Nelson and Porter, 2011), so we adjusted the associated collection date in our dataset.) The Royal Botanic Garden Edinburgh (E) houses an 1827-dated specimen of *S. galapagense* collected by John Scouler (in January 1825; Scouler, 1905) that appears on the same contemporary specimen sheet as an 1837 collection of *Solanum lycopersicum* (collector unknown). Altogether we found 19 pre-1900 specimen records of Galápagos endemic tomatoes, 12 *S. galapagense* and six *S. cheesmaniae* (Table S1). For four 19^th^ century collections we could not confirm or infer a specific island collection site within the archipelago (including Scouler’s 1825 collection), so these specimens were not included in the unique Location-Year-Species dataset used for downstream analyses (see below; Table S2).

Finally, because we were specifically interested in understanding the timing of introduction of *S. pimpinellifolium* on the Galápagos, we individually cross-checked all apparent herbarium or other collection records of this species made prior to 2000. Of these, two were misnamed specimens of *S. cheesmaniae*, reflecting past taxonomic uncertainty in this group (Darwin et al., 2003). The remaining pre-2000 records consisted of two TGRC records from each of 1985 and 1991 (LA2857 and LA3123, respectively), two 1956 specimens in the DAV herbarium, and one 1905 specimen at the California Academy/CAS (Figure 1). The TGRC records have associated germplasm which clearly confirms their identification as *S. pimpinellifolium*. The older three records are further discussed in the *Results*.

#### B. Data coding from herbarium and other collection records

Our final dataset of 410 unique historical records includes specimens from 18 herbaria, as well as TGRC and CDS collection/observation records, in addition to 44 contemporary collection records made by us during three recent field surveys (2018-2021) (Table S2). Fifteen of these records are of proposed or confirmed hybrids (*S. pimpinellifolium* x *S. cheesmaniae*).

For each unique record, our dataset at minimum includes information on species identity, collection year, herbarium/institution, island, and latitude/longitude of the collection location. In many cases latitude and longitude coordinates were not recorded with an original herbarium specimen or collection record, but had been subsequently imputed by collections managers based on field collection notes and/or on the passport label. We crosschecked these imputed latitude/longitude data against island location information to ensure that each collection record was located on the correct island. In a small number of cases, the imputed geolocation was inconsistent with the collection record (for example, the geolocation was placed in the ocean). In these cases (N=42), when passport label data were available for the specimen, we used these field notes and associated information to adjust the Latitude and Longitude of the collection location, using Google Maps. Our dataset notes all instances where we adjusted the existing latitude/longitude information for a record. For 34 records where no geographical coordinates were provided with a database record but where the passport label or collection notes included annotations about the collection location, we used these field notes to impute the Latitude and Longitude of the collection location in Google Maps. Our dataset notes all instances where we imputed the latitude/longitude information for a record that was missing these data. Otherwise, our analyses used geolocation data as reported in the associated herbarium database and/or published collection record.

#### C. Bioclimate and environmental data

Based on geolocation data for each collection record, we obtained matching temperature and precipitation data for 19 bioclimatic variables at 30 arcseconds from WorldClim (https://www.worldclim.org/data/worldclim21.html). We used the R packages *raster* (Hijmans 2024), and *sp* (Pebesma and Bivand, 2005; Bivand et al., 2013) to create a target raster of the Galápagos, conduct bilinear resampling of bioclimatic data for this target raster, and extract bioclimatic data for each sample’s coordinates. Eight records were not automatically matched with bioclimatic data because they occurred close to coastlines on smaller islands, so we manually completed each record using the bioclimatic data of the closest occurrence record, which was always within 15 arcseconds (<0.5km) on the same small island. (This interpolation is noted for each of these records; Table S2. Six of these cases were for *S. galapagense* records on Bartolomé, an island whose bioclimatic data did not vary across any collection site.) Data on individual island size (land area, in hectares) was taken from Phillips et al. (2012).

### Analyses

#### A. Bioclimatic envelope analysis

We ran a principal components analysis (PCA) on all samples (except for the 15 hybrid *S. cheesmaniae* x *S. pimpinellifolium* records) for 19 bioclimatic variables (Table S3). To assess the contribution of island and species differences to each of the first 5 principal components (PCs), we conducted 2-way ANOVAs (by island and species identity) on each PC, followed by post-hoc Tukey’s HSD Test of pairwise species differences. We also compared the variance of PCs 1 through 5 among species, with Levene’s Test in the R package *car* (Fox & Weisberg 2019). As follow-up analyses, we conducted several parallel comparisons of individual bioclimate variables, focusing on a subset that loaded most strongly on the first three PC axes.

#### B. Species distribution modelling and niche overlap

We used species distribution models (SDMs) to identify habitable regions for each species, based on climatic variables. We created SDMs using Maxent, via the R package *dismo* (Hijmans et al., 2024), following the procedure described by Feng, Gebresenbet & Walker (2017) (github.com/shandongfx/workshop_maxent_R). Because our primary aim was exploratory analysis of bioclimatic envelopes, rather than hypothesis testing, we included all sample coordinates (presence sites) for each species’ SDM. Background sampling was designed to account for the fact that field-observed occurrence data can be biased by variation in the accessibility of particular sites or habitats (e.g. Daru et al., 2017). (For instance, some locations may harbor no sample records because no observer travelled to that region, rather than because tomato populations are truly absent.) We therefore restricted background sampling to an area within 0.05 degrees (5 km) of the northern-, southern-, western-, and easternmost presence sites within each island for a given species. Background sites were randomly selected from 10% of the total number of cells within this area. All nineteen bioclimatic variables were used for the environmental layer. We repeated model generation 10 times per species and averaged the prediction rasters. We delimited habitat suitability for each species under three widely used thresholds for SDMs: minimum training presence, tenth percentile training presence, and maximum training sensitivity plus specificity. We present results based on the latter two thresholds only, selected to bracket a range of projections based biological and geographical knowledge of the focal species. We also used SDMs to identify the degree of ecological coincidence/overlap between pairs of each species, quantified by calculating Schoener’s Distance/D (Schoener 1968), with the R package *dismo* (Hijmans et al. 2023).

#### C. Visualizing temporal and spatial patterns

We visualized temporal and spatial patterns of sampling, the change in the sampling abundance per species over time, and outputs from SDMs, with R packages *dismo* (Hijmans et al. 2023), *dpylr* (Wickham et al. 2023), *raster* (Hijmans 2024), *sf* (Pebesma, 2018; Pebesma and Bivand, 2023), *sp* (Pebesma and Bivand 2005, Bivand et al. 2013), *terra* (Hijmans, 2024), *tidyterra* (Hernangómez, 2023), and *ggplot2* (Wickham 2016).

## Results

### 1. Historical records bracket the island distribution of Galápagos endemics and the timing of emergence of the introduced species

Our dataset identified 410 unique date and location records for *S. cheesmaniae* (N=169), *S. galapagense* (N=144), *and S. pimpinellifolium* (N=82) (Figure 1), as well as 15 *S. cheesemaniae* X *pimpinellifolium* hybrid records (Table S2). These records collectively span 226 years, from 1795 to 2021 (Figure S1). The endemics *S. cheesmaniae* and *S. galapagense* have been identified on 10 and 13 unique islands, respectively (Table S4). In their taxonomic revision of these species, Darwin et al. (2003) reported similar island distributions, based on S. Darwin’s field surveys in 2000/2001, and on 102 herbarium specimens (43 and 59 of *S. cheesmaniae* and *S. galapagense*, respectively). As far as we can ascertain, our dataset includes all herbarium specimens referenced in this 2003 taxonomic revision (with the exception of one 1983 *S. galapagense* record from Darwin Island, which could not be relocated in the available CDS records). Most of the additional 200+ geolocated records for the two endemic species in our dataset are drawn from herbarium records in the California Academy (CAS) and the U. C. Davis Center for Plant Diversity Herbarium (DAV), TGRC and post-2000 CDS collection records, and our own 2018-2021 Galápagos location records (Table S2).

Together these data indicate evident differences in the biogeographical distribution of the two endemic species across islands of the archipelago (Figure 2, side panels). In particular, *S. cheesmaniae* has only been recorded on islands >1800 hectares; Pinzón, at 18.15km^2^, is the smallest island with an historical collection record for this species (Table S4). In comparison, *S. galapagense* is found both on large (Isabela, Fernandina, and Santiago) and much smaller islands, including at least three islands or islets under 100 hectares (1km^2^). Interestingly, we find no verified records of *S. galapagense* on Santa Cruz, San Cristóbal, or two other islands (Santa Fé and Baltra) that are >2500 hectares (Table S4). These broad distributional differences suggest that island identity could have strong effects on species differences in bioclimate variation, as addressed further below.

**Figure 2.**
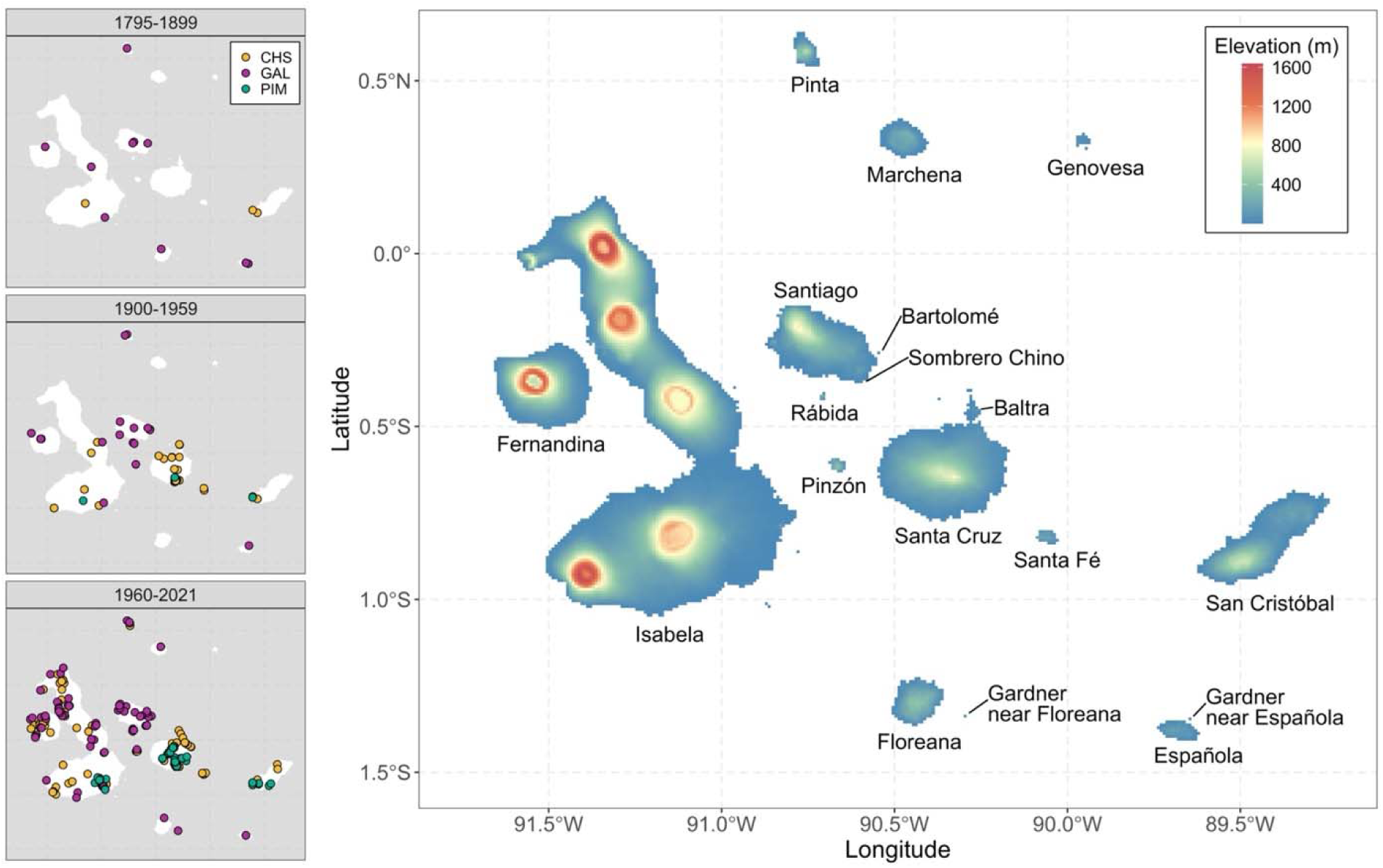
Islands in the Galápagos archipelago. Main: Map with landmass elevation (in metres). **Side panels:** Temporal and island distribution of 395 historical records of *S. cheesmaniae* (CHS, yellow), *S. galapagense* (GAL, magenta), and *S. pimpinellifolium* (PIM, green) during three time periods (top to bottom) since the late 18^th^ Century.

In comparison, introduced *S. pimpinellifolium* is only historically recorded from three islands, all human-occupied, with the majority of records on Santa Cruz (N=54), followed by San Cristóbal (N=17), and then Isabela (N=11) (Figure 2). In agreement with Darwin et al. (2003), we infer that the oldest unambiguous Galápagos collection of this species was made in 1985 on Isabela; this collection has a current, verified, germplasm collection (accession LA2857 in the TGRC). In contrast, three older Galápagos collections remain ambiguous. Two of these, collected 1956 on each of San Cristóbal and Santa Cruz (from records in the U.C. Davis Center for Plant Diversity Herbarium), do not yet have digitized images. However, digitized collection notes (Table S2) indicate one or both might instead be *S. cheesmaniae* specimens. The San Cristóbal specimen notes specify: “Only one plant seen. This form may approach *L. cheesemanii*. Need comparison in culture for better determination.” Also consistent with misattribution, there are no further records of *S. pimpinellifolium* on San Cristóbal until 2000 (Darwin et al., 2003). In comparison, the 1956 Santa Cruz record is more equivocal. Its specimen notes state that it “resembles the typical Santa Cruz type in most respects”, and *S. cheesmaniae* is now recognized to have an intraspecific variant that was historically referred to as a ‘Santa Cruz’ variant of *S. pimpinellifolium* (Rick 1956; Darwin et al. 2003). On the other hand, we previously inferred that one mainland Ecuador accession of *S. pimpinellifolium*, collected in 1956, was the result of back-migration from the Galápagos (Gibson et al. 2021; see ‘Study System’); its collection remarks specifically note phenotypic similarities to “Santa Cruz pimpinellifolium”. This implies a presence on the Galápagos since at least 1956, and a specific connection to Santa Cruz Island.

The digitized image of the 1905 specimen from Isabela (Figure 1) indicates this collection is also taxonomically ambiguous, as noted on the specimen itself (by S. Knapp; NHM) and in Darwin et al. (2003). However, in this case leaf and flower morphology more strongly suggest it might be a *S. pimpinellifolium* collection or possibly even a variety of domesticated tomato, rather than misclassified *S. cheesmaniae*. If it is *S. pimpinellifolium*, the leaf morphology of this 1905 specimen differs radically from our recent collections of this species (Figure 1), including specifically on Isabela. Specimen morphology also departs from collections made by S. Darwin in 2000 (see Figure 4 in Darwin et al., 2003). Although leaf morphology can be influenced by developmental environment (i.e. can be plastic), these evident leaf differences suggest that the 1905 specimen might represent a *S. pimpinellifolium* (or related species) introduction event that is independent of the three we have already identified with genomic data (Gibson et al. 2021), including the introduction that gave rise to most contemporary populations observed on the islands. This hypothesis—and the species identity of the 1905 and two 1956 specimens—would be testable with future genotype data from these herbarium specimens.

Finally, we note that the CDS has records of two additional, recent, collections of *S. pimpinellifolium*—one each from Floreana and Española islands (in 2002 and 2021, respectively; Table S4). Because we currently lack image or germplasm confirmation of these novel observations, they were not included in our quantitative analyses for *S. pimpinellifolium*; however, we further address their possible significance in the *Discussion*.

#### 2. Species differ significantly in bioclimate envelopes, with large inputs from regional island-to-island variation

Both principal component analyses (Figure 3, Table S5, S6), and follow-up univariate analyses (Table S7) indicate that our samples vary broadly in environmental features of their occurrence locations, especially on important axes of temperature, precipitation, and daily and seasonal climate variability. Together, the first five principal components (PCs) explained >98% of the variation in our dataset (Table S5). PC1, which captured 41.8% of the variance, loaded consistently strongly for temperature variables (Table S6). PC2 (33.2% of variance) loaded most strongly for precipitation variables (+), and to a lesser degree isothermality (-), and mean diurnal range (-) (Table S6). PC3 (9.8%) loads strongly on temperature seasonality (-), mean diurnal range (+), and isothermality (+) (Table S6). PC4 and 5 (together explaining 13.9% of the variance) most strongly load on variation or extremes in temperature and precipitation (specifically, annual temperature range (both PCs), mean diurnal temperature range and precipitation seasonality (PC4), and precipitation in the coldest quarter (PC5); Table S6).

**Figure 3.**
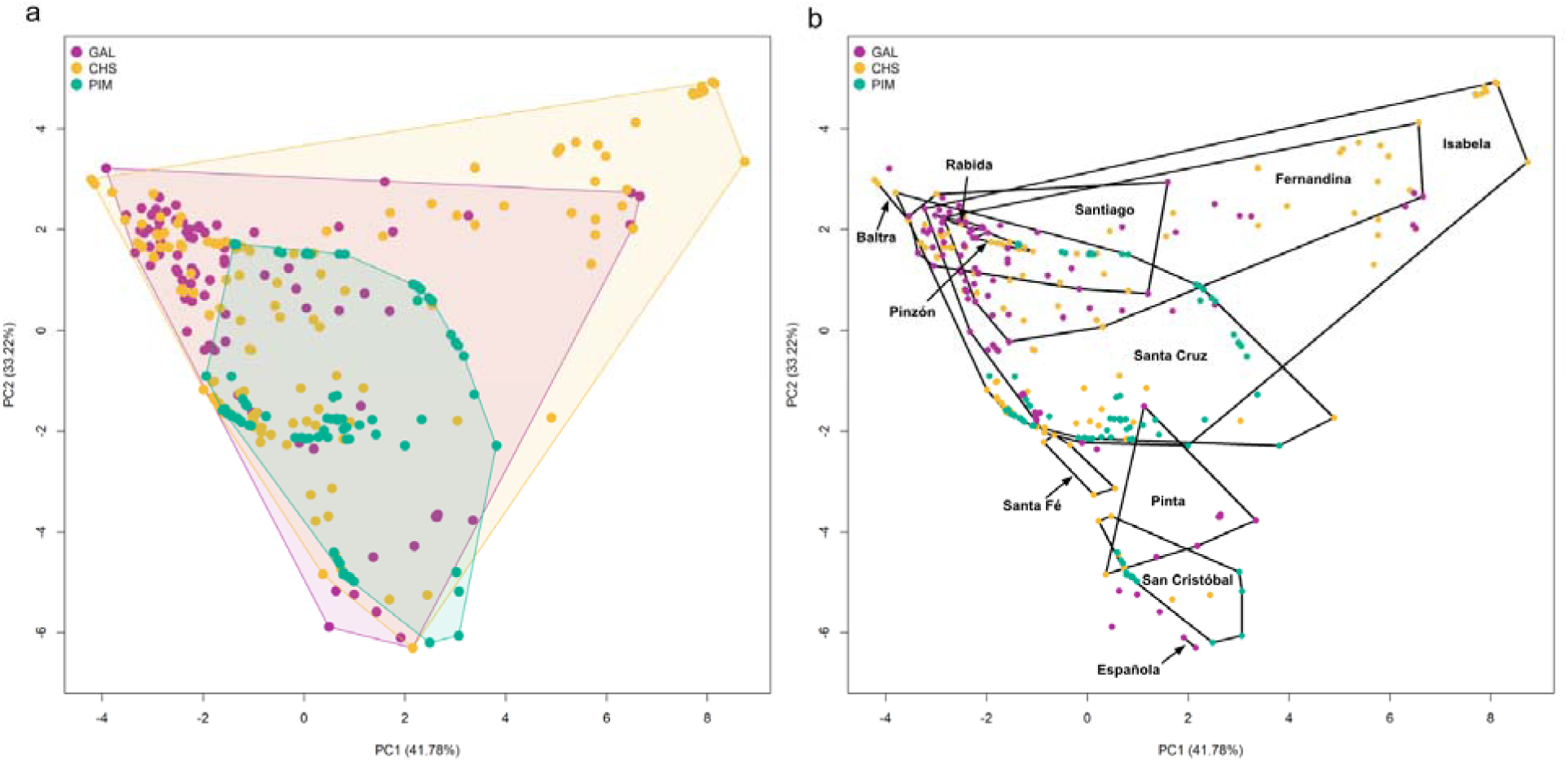
Bioclimate envelopes of the first two principal components generated from 19 bioclimatic variables, evaluated for 395 records of wild tomatoes on the Galápagos. Yellow = *S. cheesmaniae (CHS)*, Magenta = *S. galapagense (GAL)*, Green = *S. pimpinellifolium (PIM)*. **A. (left panel)** Colored polygons enclose the maximum extent of all records for each species. **B. (right panel)** Black polygons bracket the maximum extent of records from individual islands, as labelled. Only islands with 2 or more records bearing different PC values are delimited with polygons. The predominant loadings on each axis are described in the main text (see *Results*) and Table S6.

**Figure 4.**
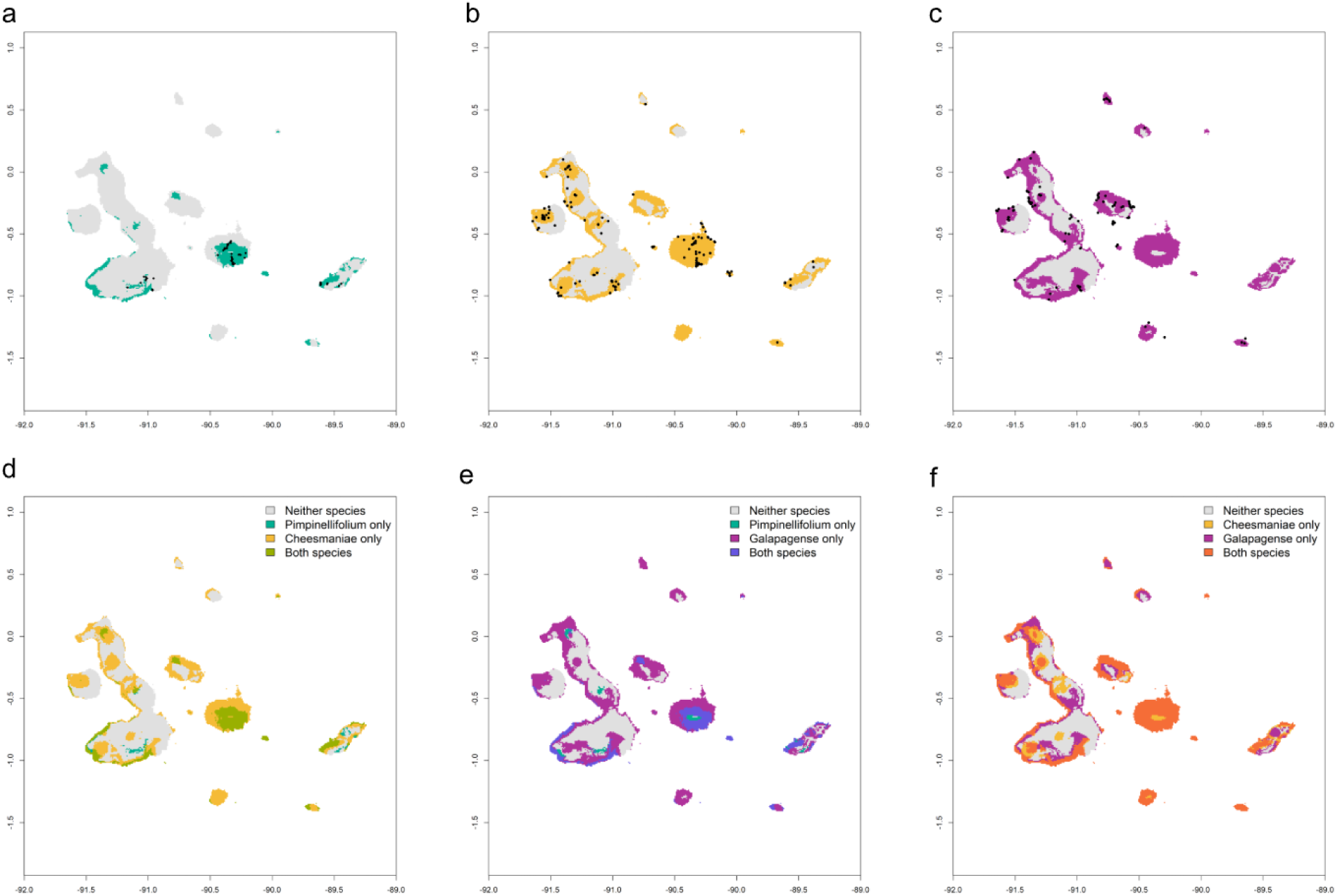
Species distribution models for three wild tomato species on the Galápagos, using 10^th^ percentile training presence threshold. **Upper panels:** Predicted suitable habitat for (a) *S. pimpinellifolium*, (b) S*. cheesmaniae*, and (c) *S. galapagense*. Dark points indicate actual occurrence records, shaded areas indicate predicted suitable habitat. **Lower Panels:** Predicted niche overlap between (d) *S. pimpinellifolium* & S*. cheesmaniae*, (e) *S. pimpinellifolium* & *S. galapagense*, and (f) S*. cheesmaniae* & *S. galapagense.* Schoener’s D estimates for niche overlap are given in Table S9.

Interestingly, our PCA suggests that island-associated bioclimatic differences (Figure 3B) are at least as pronounced as archipelago-wide species differences (Figure 3A). This inference is supported by strong significant island effects on PCs1 through 5 in two-way ANOVAs that include both island and species as main effects (Table S7). Island is a stronger predictor of all five PCs, although species is also a strong predictor of variation in PCs 1, 2, and (especially) 3 (Table S7), independent of island. In post-hoc tests, PC1 (∼ temperature) differentiates *S. galapagense* from *S. cheesmaniae* and *S. pimpinellifolium* (Table S7); PC2 (∼ precipitation) differentiates *S. cheesmaniae* from the other two species; and PC3 (that captures elements of daily and seasonal climate variation) strongly differentiates introduced *S. pimpinellifolium* from the two endemics (Table S7). We found similar results for individual bioclimate variables that strong load on each of these three PCs (Table S8). Nonetheless, the most prominent bioclimatic axes in our dataset are clearly strongly influenced by regional biogeographic (island-to-island) variation, as well as more local environmental differences within islands.

#### 3. Models of habitat suitability built from historical occurrence records differentiate endemic species preferences for alternative island climate zones

Our species distribution models (SDMs) also align with these PCA results, identifying significant species differences in projected habitat suitability, including across islands. As is expected, a more permissive threshold generated larger projected areas of suitable habitat that captured more of the known occurrence locations for all three species (especially for *S. cheesmaniae*) (Figure 4) compared to a more stringent threshold (Figure S2). Nonetheless, projections under both thresholds revealed several general patterns. Most notably, inferred suitable habitat for each species broadly coincides with geographical boundaries of recognized climate zones within and among specific islands (especially ‘arid’, ‘transition’, and ‘humid’ zones; Tye and Francisco-Ortega, 2011; and Figure 3 in Trueman and d’Ozouville, 2010). For instance, the projected suitable habitat for *S. galapagense* (Figure 4c) encompasses the geographical distribution of ‘arid’ zones across the archipelago, consistent with this species frequent occurrence in arid lowland habitats, including on bare volcanic rock (e.g., Figure 1e). *S. galapagense*’s projected suitability also extends well into the ‘transition’ climate zone on most islands—including substantial projected areas on islands where this species does not occur (e.g. Santa Cruz, Santa Fé, San Cristóbal: Figure 4c)—likely because it occurs in transition zones on Isabela, Santiago, and Pinta. Conversely, habitat projections exclude *S. galapagense* from higher elevation, humid ecosystem, locations (Figure 4c) which occur exclusively on larger islands. Suitable habitat also includes some high calderas on Isabela and Fernandina (Figure 4c), which are also locally arid and often have recent bare lava (e.g. Figure 3 in Trueman and d’Ozouville, 2010).

*S. cheesmaniae*’s suitable habitat projection coincides substantially with *S. galapagense*’s (Figure 4f), consistent with *S. cheesmaniae* also occurring in transition ecosystems. Estimated niche overlap is also high between the two endemic species (Schoener’s D_CHS,GAL_ = 0.7032 – 0.6055, depending upon the threshold; Table S9). However, unlike *S. galapagense*, *S. cheesmaniae*’s projections extend into higher elevation humid ecosystems, especially on Santa Cruz and Isabela (Figure 4b,f, and Figure S2b,f); they also avoid specific areas of dry coastal lowland, especially on north and eastern coasts of Isabela, and on the smaller lower-lying islands. Nonetheless, compared to *S. galapagense*, *S. cheesmaniae*’s projections are less clearly aligned with simple climatic zonation, and appear to be more sensitive to model decisions. For instance, at the less stringent threshold, projected suitable habitat includes the small islands Genovesa and Marchena (Figure 4b) that are entirely arid lowlands or bare volcanic rock (Figure 2, main map). In comparison, a more stringent threshold nearly or completely excludes these locations as suitable *S. cheesmaniae* habitat (Figure S2b), but poorly captures known occurrence locations of this species, especially on Isabela and San Cristóbal (see Figure S2b). It is possible that three singular occurrence records of this species, on each of Santiago, Pinta, and Española, are hampering more stable model projections, especially at the margins of this species habitat preferences. This heterogeneity might also reflect real biological variation across populations or genotypes of this species.

#### 4. Bioclimate data and species distribution models for S. pimpinellifolium indicate its potential impacts, and environmental limits, on the archipelago

In comparison to the two endemics, introduced *S. pimpinellifolium* has the narrowest range of projected habitat suitability (Figure 4a), consistent with its confirmed presence on only three islands (Figure 2) and its comparatively narrow bioclimatic envelope (Figure 3). *S. pimpinellifolium* also has the fewest occurrence records and—given its recent introduction—is unlikely to be at equilibrium with respect to exploring niche space within the Galápagos. As a result, bioclimatic envelopes and the presence-only Maxent model—both built solely from current collections—may not capture *S. pimpinellifolium*’s full range of climate preferences or its expansion potential. Nonetheless, even this exploratory analysis allows us to draw several provisional empirical inferences.

First, *S. pimpinellifolium* suitable habitat is inferred to be concentrated on larger islands with higher elevation climate zones. Projected suitability coincides with ‘humid’ highland ecosystems on several larger islands (Figure 4a), including in locations where both endemic species are projected to be absent (on Isabela and San Cristóbal especially) (Figure 4d,e). Regardless of threshold, models also place *S. pimpinellifolium* in locations where it is not yet found, including on Fernandina, Santiago, Española, and (especially) Santa Fé, as well as remote locations on western and northern Isabela (Figure 4, Figure S2).

Second, *S. pimpinellifolium*’s projected niche overlap is consistently greater with endemic *S. cheesmaniae*, compared to *S. galapagense* (10^th^ percentile threshold: D_CHS,PIM_ = 0.2673056, D_GAL,PIM_ = 0.217043; even larger differences were observed for the more stringent threshold; Table S9). This overlap largely coincides with shared humid and transition zones on Isabela, Santa Cruz, and San Cristóbal, and to some extent Santiago (Figure 4d). It also encompasses geographical locations where these species are known to currently cooccur and hybridize on Santa Cruz (Table S2) specifically in the transition zone, adjacent to human disturbed sites including roads, mines, the municipal dump, and associated waste areas (Darwin 2009, Nuez et al. 2022, Gibson et al., 2020). Although projections also appear to show large areas of overlap between *S. pimpinellifolium* and *S. galapagense* on San Cristóbal, Santa Cruz, and Santa Fé (Figure 4e), these are unlikely to represent true threats to the endemic species, as *S. galapagense* is not known to occur on any of these islands, either currently or in the past.

Third, known occurrence locations and projected habitat suitability suggest ecological constraints on *S. pimpinellifolium*’s distribution. This species’ projected suitable habitat is sparse or absent on smaller islands that are dominated by arid conditions (Figure 3, Figure S2). Within Santa Cruz—the island on which this species has had the greatest opportunity to spread—*S. pimpinellifolium* also appears to experience a distributional boundary imposed by precipitation (Figure 4a, Figure S4). Moreover, bioclimatic data from known hybrid sites on Santa Cruz (Table S2) also suggest these hybrids occur up to the edge, but not beyond, the lower precipitation range of their introduced parent species (Figure S4); no such clear boundary is evident for other bioclimatic axes, such as temperature. These observations hint at ecological constraints preventing the spread of *S. pimpinellifolium* and its hybrid offspring into more arid locations, consistent with physiological limits on tolerance for low moisture/low precipitation field conditions in these introduced genotypes.

## Discussion

The characterization of species distributions over space and time benefits uniquely from historical scientific records. Early 20^th^ century assessments of Galápagos tomato species drew substantially from the Hopkins-Stanford (1898-1899), and California Academy (1905-1906) expeditions (Robinson, 1902; Stewart, 2011). Compared to other early scientific collecting in the archipelago (including the H. M. S. Beagle visit in 1835), each of these extraordinary scientific efforts lasted for multiple months and was relatively systematic in visiting the majority of islands of substantial size (Table S4). Nonetheless, our data indicate that the subsequent 120 years of collections have significantly expanded and deepened the distribution and occurrence information on these species. Our goal in this study was to aggregate all current digitized records of wild tomato species on the Galápagos, to generate an enriched historical record of species presence, and to quantitatively evaluate species differences, risks, and threats. These data enable us to draw more robust and specific inferences about the past biological and demographic history of these species, including over recent timescales that were substantially affected by human impacts. They also allow us to make tentative qualitative and quantitative projections for each species into the future.

### Past and present dynamics of endemic and introduced wild Galápagos tomato species

First, more than two centuries of occurrence records enabled us to confirm qualitative and quantitative differences in species’ bioclimate characteristics. Compared to introduced populations of *S. pimpinellifolium,* both endemic species occur across a broader range of island environmental conditions (Figure 3, Figure S3) especially extending into more arid and warmer geolocations (Table S8). These endemic distributions are consistent with a longer-term history of adaptation to the predominant climatic conditions on the archipelago, where the arid zone represents >80% of the total land area (Camacho-Escobar et al., 2021) and may have been even greater during past glacial/ice-age cycles (Jackson 1993). Nonetheless, the two endemics also clearly differ in their climatic preferences, including island-to-island differences, reflecting important physiological differences between them. Of the >230 islands, islets, and rocks enclosed within the Galápagos Marine Reserve (Escobar-Camacho et al., 2021), only those whose elevation exceeds ∼200m experience more permanent dense fog (Porter, 1979). *S. cheesmaniae*’s apparent restriction to larger, higher islands is consistent with limits on this species’ tolerance of persistent aridity, periodic severe droughts, and the lack of soils that characterize smaller, lower elevation islands (Tye and Francisco-Ortega, 2011; Stoops, 2014). In comparison, *S. galapagense* is known to have multiple traits associated with sustained drought tolerance (e.g., Fenstemaker et al., 2022), perhaps reflecting adaptive responses to more extreme aridity over evolutionary timescales, compared to its sister species.

Second, in tandem with inferred environmental preferences, our collections data also allows us to interpret current endemic distributions in the light of historical geolocations. For instance, records from across two centuries indicate that *S. galapagense* was historically absent on the north-eastern islands of Santa Cruz, Baltra, Santa Fé, San Cristóbal, and Genovesa, rather than having been recently extirpated from these locations. Because these are islands where we also infer the presence of substantial suitable habitat for *S. galapagense* (Figure 4, Figure S2), its absence therefore suggests other factors, such as historical dispersal limitation, could explain the uneven distribution of this species across the archipelago. The origin and pathway(s) of island expansion are potentially discoverable with demographic reconstruction from genomic data (e.g. Nuez et al. 2004, Pailles et al., 2017), ideally in conjunction with geological information about the history of sea-level change and island connectedness over the last 0.5 MY (Perry, 1984; Norder et al., 2018).

Third, our collections data allows us to interpret the impact of recent environmental change on endemics’ historical distributions. Our dataset spans a period of time marked by the expansion of permanent human settlement and agriculture, the introduction of large herbivores and alien plant species, and a rapidly growing tourist industry. Our collections dataset reveals a heterogeneous footprint of this recent environmental change, that is most evident in collections data from human-occupied islands (Figure 1). For instance, since 1985 the human population increased from ∼6000 to >30,000, and tourism grew from ∼20,000 to >270,000 people per year (Phillips et al., 2012, Escobar-Camacho et al., 2021). Over the same period, on the two most populated and heavily trafficked islands—Santa Cruz and San Cristóbal—*S. cheesmaniae* collection records vanish from locations adjacent to human settlements, as well as almost entirely from the humid zone generally (Table S2, Figure 2). On Isabela, new *S. cheesmaniae* records expand over this same period, but only remotely from human-impacted locations (Figure 2). Similar temporal changes are not evident in records on more remote islands, in spite of overall lower collecting intensity. For instance, although the 20^th^ century collection frequency of *S. galapagense* is consistently lower than *S. cheesmaniae* (Figure S1, Table S2), its collections are comparatively stable over this period because of records from smaller or more remote islands. On Bartolomé, for example, populations of *S. galapagense* have been collected across nearly seven decades, often at geolocations within tens of metres of each other (Table S2). Even Santiago—whose endemic plant communities were devastated by introduced herbivores (prior the systematic eradication of goats, donkeys, and pigs by the early 2000s; Phillips et al. 2012)—has consistent collections into the current century (Figure 2). Interestingly, occurrence remarks from several mid-20^th^ century records specifically note that individuals were growing in a location inaccessible to goats (Table S2; see also Nuez et al., 2004). The resilience of *S. galapagense* to these effects therefore appears to be associated with its preference for physiologically challenging, often remote and inaccessible, locations that are buffered from the most overt impacts of humans and the plants and animals that have accompanied them.

Finally, our collections dataset allows us to interpret the current distribution of the introduced *S. pimpinellifolium* in the light of unique historical records of this species. Our previous genotyping supported three separate introductions of *S. pimpinellifolium* onto the Galápagos; one older introduction from Ecuador that gave rise to most contemporary sampled populations on Santa Cruz and San Cristóbal, and two more recent introductions from Peru (onto Isabela) and mainland Ecuador (onto Santa Cruz) (Gibson et al., 2021). We do not yet know the relationship between the earliest confirmed *S. pimpinellifolium* on the Galápagos (TGRC accession LA2857; from 1985), and any of our genotyped specimens, although our Isabela collection (MG128; from 2018) was also made in the town of Puerto Villamil. Genotyping data could resolve whether the 1985 collection draws from one of the same introductions or represents a fourth independent event. Future genotyping would also resolve the status of the three earlier 20^th^ century collection records of this species, including the phenotypically unusual specimen from 1905 (Figure 1). Of these three, the 1956 specimen from Santa Cruz is potentially most interesting with respect to understanding sources of currently invasive populations. Prior genotyping implies a presence on Santa Cruz since at least 1956 (see ‘Study System’), and Santa Cruz also has the highest current abundance of *S. pimpinellifolium* (Gibson et al. 2020), with notable populations both inside and outside the higher elevation humid zone since the late 1990s (Darwin, 2009; Nuez et al., 2004). Regardless of their specific relationships, it is clear that island records of this species begin and persist in close contact with human settlements and disturbance, including every collection record made prior to 2000.

### Future projections for endemic and introduced wild Galápagos tomato species

Compared to assessing past dynamics, extrapolating into the future is clearly more tentative. Nonetheless, two general inferences suggest themselves from our analyses of historical records of these species. First, we infer more, and more acute, future conservation threats to *S. cheesmaniae*. The major projected drivers of change in terrestrial Galápagos ecosystems are climate change, unsustainable tourism and local population growth, and invasive species (synthesized in Escobar-Camacho et al., 2021). *S. cheesmaniae*’s bioclimatic distribution— specifically its apparent limitation to larger higher islands, including those with a substantial human presence—amplifies its exposure to all three drivers. Indeed, in spatial impact assessment models, projected areas of greatest future ‘exposure’ to environmental threats coincide substantially with the historical distribution and projected suitable habitat of *S. cheesmaniae* (see Figure 4 from Escobar-Camacho et al., 2021; reproduced as Figure S5). In addition to these general threats, *S. cheesmaniae* faces a specific risk from ongoing and future contact with *S. pimpinellifolium*. These species are currently in physical and reproductive contact at several sites on Santa Cruz (Gibson et al., 2020), and have a history of hybridization that extends up to 12 generations (Gibson et al., 2021). Our projections of species’ suitable habitat indicates that this contact could increase in the future, especially if the realized range of *S. pimpinellifolium* continues to expand. In contrast with *S. cheesmaniae*, future conservation threats to *S. galapagense* are harder to predict. Climate models project relatively large changes in precipitation for known locations of *S. galapagense*, including southern arid lowlands on Isabela, and in arid ecosystems on Española, Marchena, Pinta, and Pinzón (Escobar-Camacho et al., 2021); this could potentially amplify exposure to new, less arid-adapted, colonizer species. Sea level in the Galápagos is also rising—by 10cm since 1985 (Escobar-Camacho et al., 2021)—threatening the preferred habitats of species like *S. galapagense* that can be found in the littoral zone, just metres above high tide.

Second, inferences from historical records also indicate the most likely paths to future invasive spread for *S. pimpinellifolium*, as well as possible limits to this expansion. Our habitat projections from current occurrence records suggest introduced *S. pimpinellifolium* prefers the cooler and (especially) wetter conditions of humid island ecosystems, perhaps due to intrinsic physiological constraints that prevent further expansion into more arid locations. Interestingly, this apparent limitation might be due to the specific *S. pimpinellifolium* genotype(s) introduced onto the Galápagos, as mainland populations of this species extend into arid coastal regions of Ecuador and Peru (Nakazato et al. 2010; Gibson and Moyle 2020) and exhibit drought-adaptive traits (e.g. Nakazato et al. 2008). Regardless, our projections also identify suitable habitat in several locations that *S. pimpinellifolium* does not yet occupy (Figure 4, S2)—including on Fernandina, Santiago, Española, and Santa Fé (Figure 4, Figure S2)—consistent with the seven Galápagos islands that are high enough to support a humid zone (Tye and Francisco-Ortega, 2011). These suitable but unoccupied habitats suggest that dispersal limitation plays an important role in shaping *S. pimpinellifolium*’s current distribution, and also identify island locations that could plausibly support viable populations of this species if dispersal was not limited. In this regard, we note that two recent records (made by the CDS; Table S4) suggest new introductions of *S. pimpinellifolium* onto each of Floreana and Española. The Floreana record (made in 2002) is geolocated in the center of the only human settlement (Puerto Velazco Ibarra), suggesting it might have been made in a waste area, domestic garden, or even a restaurant; despite its introduced status, *S. pimpinellifolium* is routinely advertised in island restaurants as ‘Galápagos tomato’ (Darwin, 2009; and pers. obs.). The 2021 Española record is from a beach location, and landing site for tourist day trips, suggesting a role for tourist traffic in its introduction. As with pre-2000 occurrence records, we do not yet know whether these events are independent of *S. pimpinellifolium* populations elsewhere on the Galápagos. However, they reinforce our inference that every early record of this species is closely associated with human activity, and underscore that restricting further expansion is strongly contingent on continuing to limit introduction via human sources (a major goal of Galápagos National Park regulations on the movement of biological materials into protected areas; Toral-Granda et al., 2017). They also make clear that the existence of physical historical specimens—and the opportunity to use these in future genetic analyses—will be an invaluable resource for further defining this species’ island origin and spread.

## Supporting information

SupplementaryTables1through9

## Acknowledgements

This research was made possible by decades of collecting and preservation work by botanists, systematists, and collections managers. We are grateful for their efforts, and for sustained institutional support of the resulting physical collections in herbaria and germplasm banks. We especially acknowledge the assistance of S. Knapp (Natural History Museum London), the prior research insights and physical collections of S. Darwin, and the botanical collection curators at the California Academy of Science, and the U. C. Davis Center for Plant Diversity Herbarium.

Digitized images of herbarium specimens (Figure 1) were reproduced with permission via Creative Commons licenses from the associated herbarium or institution (see caption to Figure 1).

## Competing Interests

The authors declare no competing interests.

## Author contributions

The research was designed by ADK, ZLH, and LCM, who also collected the data. ADK and ZLH analyzed the data, and ADK, ZLH, and LCM interpreted the results. LCM and ADK wrote the manuscript, with assistance from ZLH.

## Data Availability

All original data generated in this work are provided in the supporting information to this paper.

## Supporting Information

### Supporting Table Captions

Table S1: Pre-1900 digitized records of wild tomato collections, and associated specimens, from the Galapagos Islands.

Table S2: Dataset of unique species/date/location herbarium and other collection records of three Galapagos wild tomatoes

Table S3: Dataset of unique species/date/location records of three Galapagos wild tomatoes: Extracted raster values for 19 bioclim variables (N=410), and first 5 principle components from PCA (N=395)

Table S4: Island Distirbutions of Galapagos wild tomatoes. A) Galapagos Islands with reported occurences of one or more wild tomato species. B) Galapagos Islands or Islets >20 hectares which contain no reports of wild tomatoes.

Table S5: Principal Component Analyses of 19 bioclimate variables across Galapagos occurrence records for three wild tomato species

Table S6: Component loadings from PCA of 19 bioclimate variables across three Galapagos tomato species occurrence records

Table S7: Variation in the first 5 PCs according to species and island: (A) two-way ANOVAs and

(B) Tukey posthoc tests of pairwise species differences

Table S8: Variation in the highest loading bioclim variable for each of PC1, 2, and 3, according to species and island: (A) two-way ANOVAs and (B) Tukey posthoc tests of pairwise species differences

Table S9: Species Distribution Models (SDMs) for two alternative thresholds of predicted suitable habitat.

### Supporting Figures

**Figure S1.**
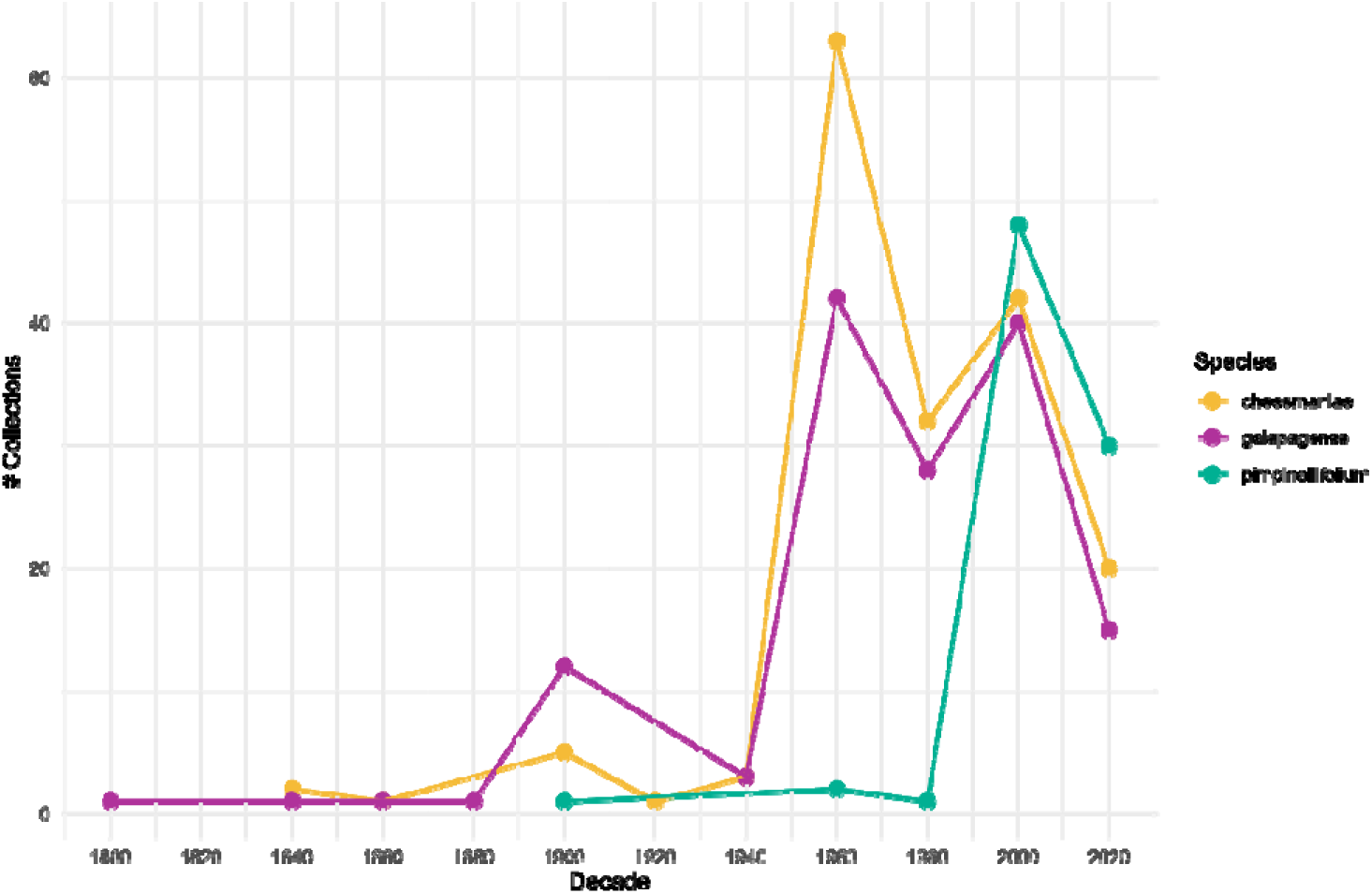
The number of herbarium specimens and other occurrence records of wild tomatoes made on the Galápagos Islands over twenty-year periods since 1800. Most collections and records have occurred since early collection expeditions, including lengthy dedicated expeditions in 1898/1899 and 1905/1906. Collections of *S. pimpinellifolium* have rapidly increased in prevalence over time, post-1980. Three pre-1985 collections require further species verification, but might represent early examples of alien introductions.

**Figure S2.**
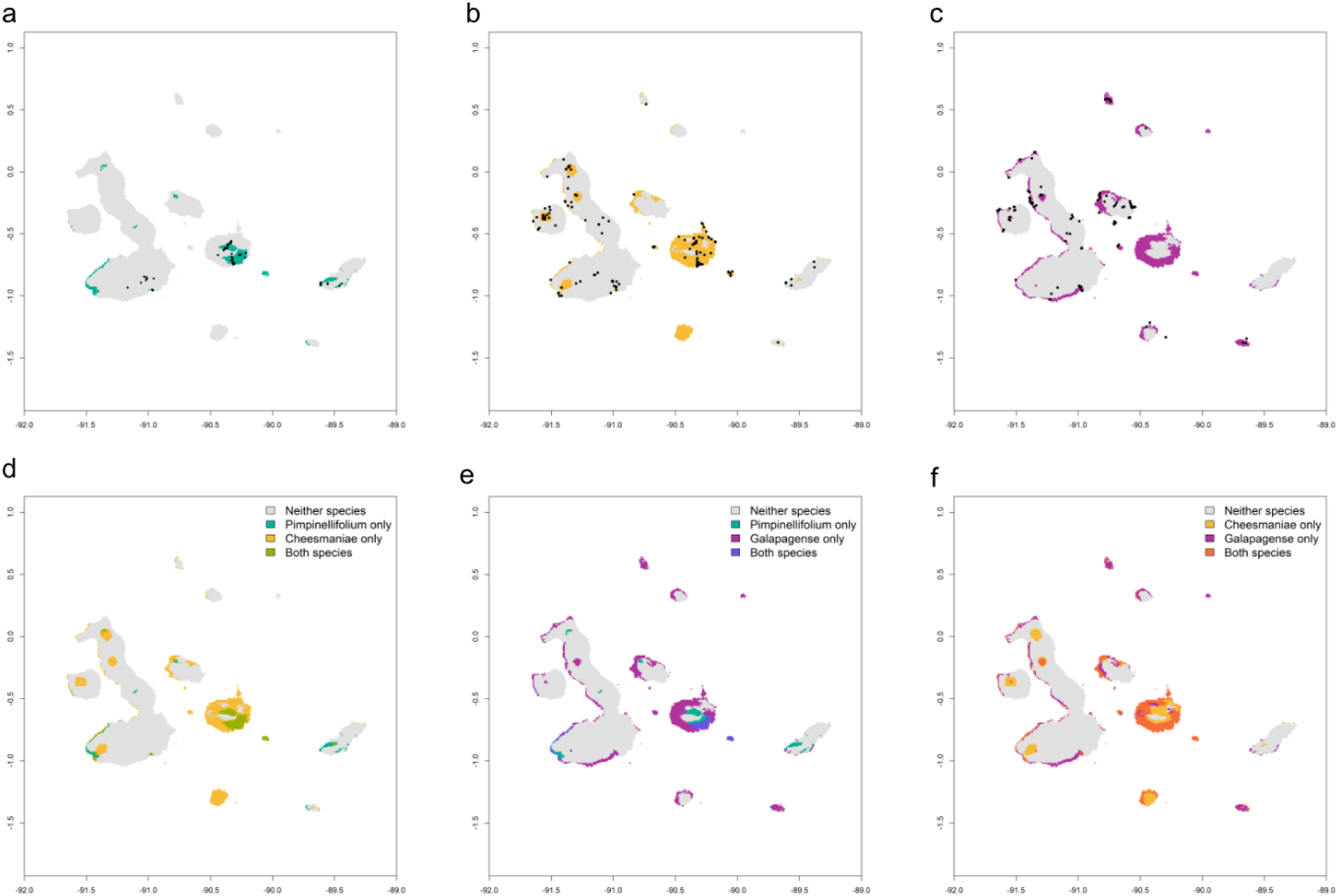
Species distribution models for three wild tomato species on the Galápagos, using maximum sensitivity and specificity threshold. **Upper panels:** Predicted suitable habitat for (a) *S. pimpinellifolium*, (b) S*. cheesmaniae*, and (c) *S. galapagense*. Dark points indicate actual occurrence records, shaded areas indicate predicted suitable habitat. **Lower Panels:** Predicted niche overlap between (d) *S. pimpinellifolium* & S*. cheesmaniae*, (e) *S. pimpinellifolium* & *S. galapagense*, and (f) S*. cheesmaniae* & *S. galapagense.* Schoener’s D estimates for niche overlap are given in Table S9.

**Figure S3.**
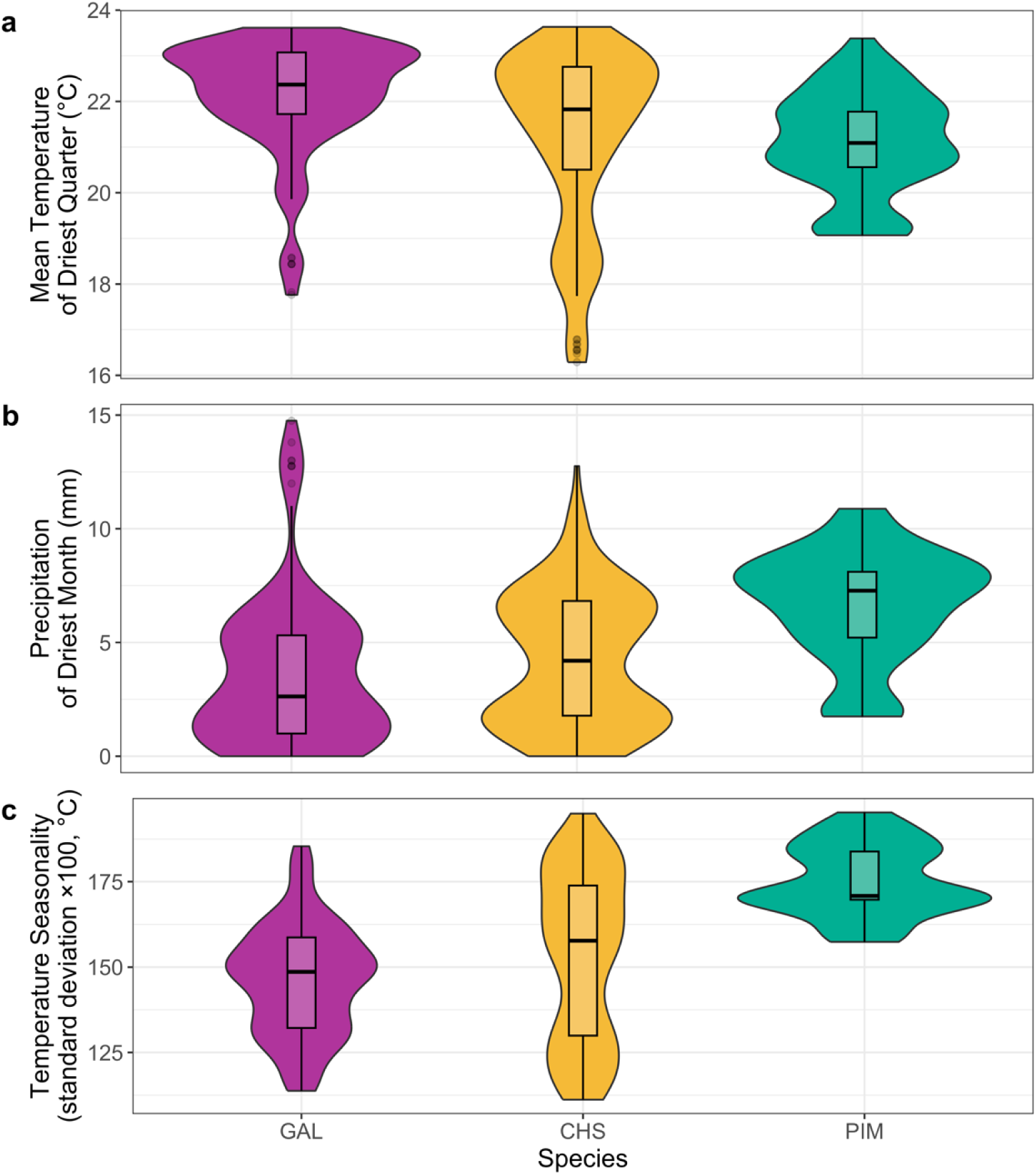
The distribution of environmental variation extracted for geolocated occurrence records of three wild tomato species on the Galápagos. GAL: *S. galapagense* (endemic), CHS: *S. cheesmaniae* (endemic), and PIM: *S. pimpinellifolium* (green). Box and whiskers describe the median, and 75% quartiles. Violin plots show density distributions. **a.** Mean temperature of driest quarter (BIO9), the highest loading factor on PC1. **b.** Precipitation of driest month (BIO14), the highest loading factor on PC2. **c.** Temperature seasonality (BIO4), the highest loading factor on PC3. Two-way ANOVAs (species and island), and species post-hoc contrasts are given in Table S8.

**Figure S4.**
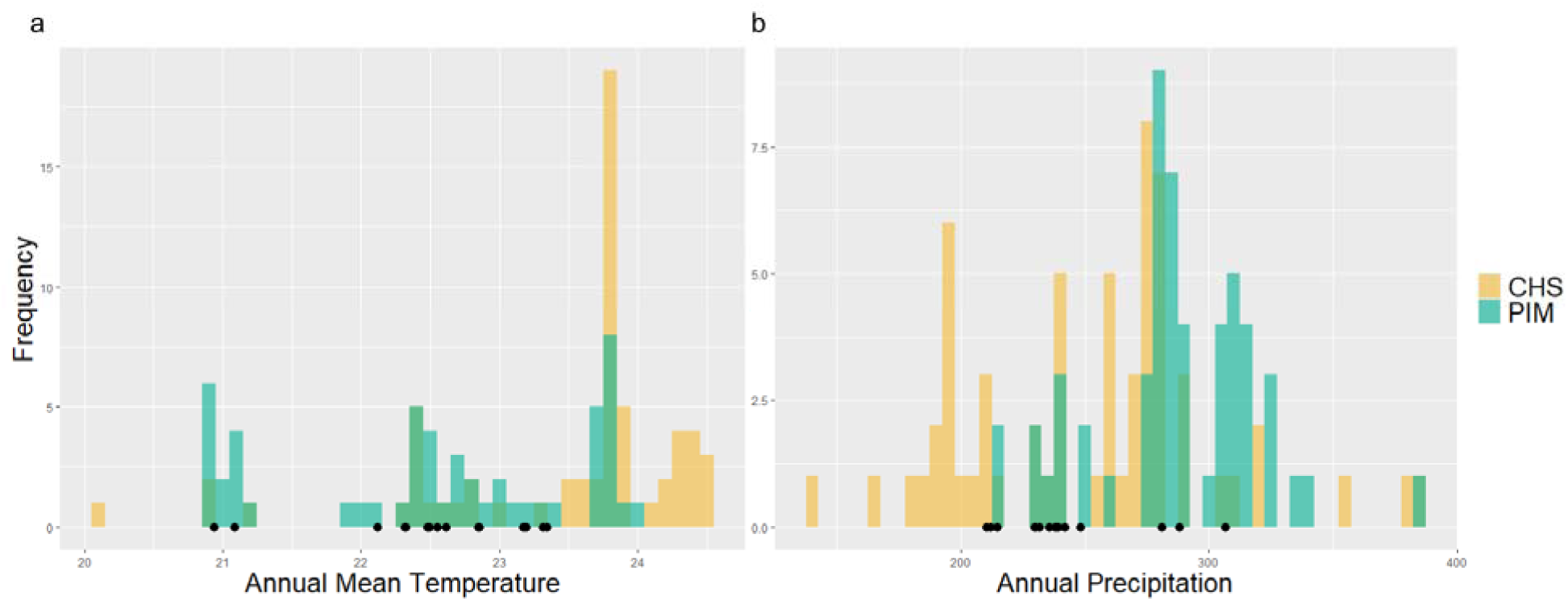
The distribution of records of *S. pimpinellifolium* (PIM; green) and *S. cheesmaniae* (CHS; yellow) on Santa Cruz, according to the (Left) annual mean temperature (increments of 0.1C) and (Right) annual precipitation (increments of 5mm) of their geolocation. Hybrids are displayed as black points.

**Figure S5.**
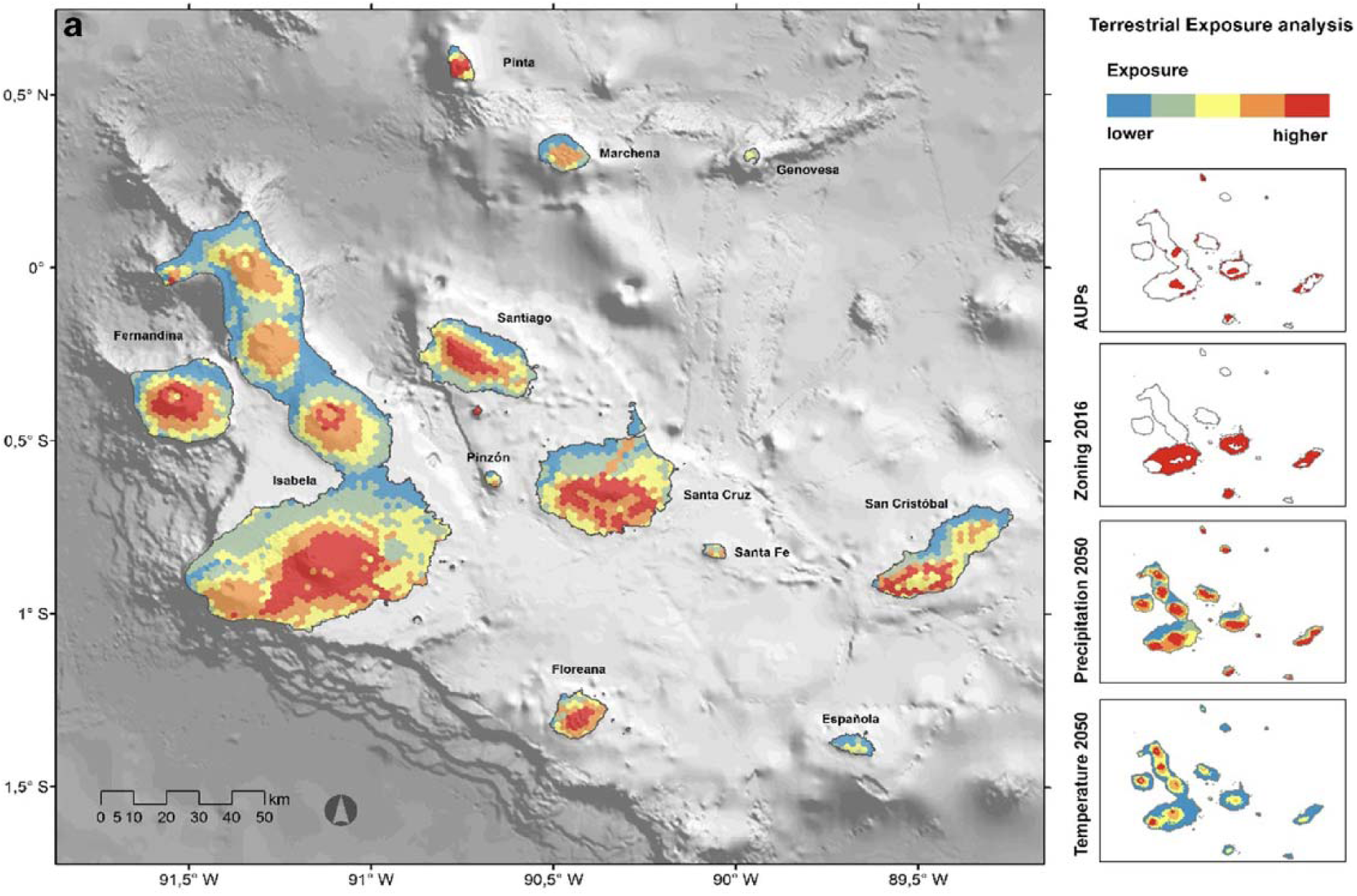
Reproduction of Figure 4A from Escobar-Camacho et al., (2021), showing their model projection of terrestrial Galápagos ecosystems exposure to environmental drivers. According to the source study, terrestrial exposure “was calculated by the admitted capacity of tourism sites (PUA), the presence of sustainable use and transition areas…, and estimated changes of precipitation and temperature for 2050” (Escobar-Camacho et al., 2021, p. 46). The spatial distribution of these four factors is displayed on left inset maps. Figure reproduced under a Creative Commons Attribution 4.0 International License (http://creativecommons.org/licenses/by/4.0/).

## Notes

### Competing Interest Statement

The authors have declared no competing interest.

